# Mass spectrometry imaging shows specific lipids signals linked to Cyp27b1 ablation in the MMTV PyMT breast cancer model

**DOI:** 10.64898/2026.08.10.743249

**Authors:** Mengdi Xing, Ethan Yang, Jiarong Li, Frédéric Fournelle, Rachel S. Pryce, Ami Grunbaum, Pierre Chaurand, Richard Kremer

## Abstract

Bioactive vitamin D (1,25-dihydoxyvitamin D or 1,25(OH)_2_D) is synthesized from its inert circulating form 25-hydroxyvitamin D (25(OH)D) by the enzyme 1-α-hydroxylase in the kidneys and in other tissues including breast. Because breast cancer is associated with changes in intra-tumoral lipid composition and vitamin D is known to affect lipid metabolism, we investigated the potential role of tumor-produced 1,25(OH)_2_D on lipid profile expression during breast tumor progression. For that purpose, we used the MMTV-PyMT mouse model which mimics the four phases of tumor progression seen in human breast cancer (hyperplasia, adenoma/mammary intraepithelial neoplasia (MIN), early carcinoma and late carcinoma). In previous studies we showed that conditional ablation of the gene encoding 1-α-hydroxylase (*Cyp27b1*), specifically in the mammary epithelium of this MMTV-PyMT mouse model, resulted in enhanced spontaneous tumor initiation and progression. In the present study, we used mass spectrometry imaging to compare lipid composition in the mammary glands of *Cyp27b1* ablated and non-ablated MMTV-PyMT mice. In non-ablated control animals, we observed changes to specific lipid signals linked to stages of tumor progression. In particular, several discriminatory lipid signals were significantly up regulated throughout tumor progression. In ablated mice, absence of *Cyp27b1* in the mammary epithelium was accompanied by different lipid signals in hyperplastic lesions. Several lipid signals were exclusively detected in non-ablated tumors but absent in hyperplasia. Our findings suggest that the tumor-produced 1,25(OH)_2_D known to play a key role in mammary tumor progression is mechanistically related to early changes in lipid composition seen prior to the development of hyperplasia.

## Introduction

Vitamin D is a member of a family of steroid hormones with pleiotropic roles in calcium and phosphate intestinal absorption, bone health maintenance, muscle and immune system function, as well as reduction of insulin resistance and prevention of pathologies such as cardiovascular disease and cancer [1–7]. Humans can obtain vitamin D from their diet, but the largest source is synthesis in skin exposed to sunlight where the precursor 7-dehydroxycholesterol is transformed into cholecalciferol. The liver then synthesizes the 25-hydroxyvitamin D (25(OH)D) prohormone through the action of cytochrome P450 [8]. 25(OH)D is the circulating form but is biologically inert and must be activated to the bioactive form 1,25-dihydroxyvitamin D (1,25(OH)_2_D) through hydroxylation by the 1-α-hydroxylase encoded by the *CYP27b1* gene present in both renal and extra-renal tissues. In kidneys, CYP27B1 is regulated by plasma calcium, phosphorus and parathyroid hormone, and the active 1,25(OH)_2_D is released to and exerts calcemic effects. Contrastingly, in extra-renal tissues, the activity of CYP27B1 is substrate-dependent and produces active 1,25(OH)_2_D that is not normally released into circulation [9], and has only local effects [10]. To exert its activity, 1,25(OH)_2_D binds the vitamin D receptor (VDR), which heterodimerizes with the retinoid X receptor (RXR) and interacts with vitamin D-responsive DNA elements, thereby regulating the expression of hundreds of genes [11].

While the classical action of vitamin D on bone mineralization as well as on calcium and phosphorus homeostasis is well-known, its non-classical involvement in cancer is a more recent but crucial observation [12]. 1,25(OH)_2_D displays antiproliferative effects in several cancers including breast [6,13–22] where it decreases *in vitro* invasion and angiogenesis [23–26]. There is universal expression of CYP27B1 and of the VDR in normal mammary glands, breast tumors and adjacent non-cancerous tissues. However, breast cancer patients have significantly reduced expression of CYP27B1 in adjacent non-cancerous tissue compared to healthy controls [27]. Expanding on the well-known inverse association between circulating 25(OH)D levels and breast cancer growth [6,7,28–30], we have previously demonstrated that tumor-synthesised 1,25(OH)_2_D delays tumor progression [6] in a mouse model. The mouse mammary cancer model MMTV-PyMT (polyoma virus middle T antigen under the control of the mouse mammary tumor virus promoter) mimics the 4 stages of breast cancer progression (hyperplasia, adenoma/mammary intraepithelial neoplasia (MIN), early carcinoma and late carcinoma) followed by metastasis to the lungs [31]. In a subsequent study, the *Cyp27b1* gene was ablated specifically in the mammary epithelium of MMTV-PyMT mice, reducing local 1,25(OH)_2_D levels and accelerating spontaneous mammary tumor development [5]. While some details of the tumor inhibition mechanisms through tumor-produced 1,25(OH)_2_D remain unresolved, it is clear that the 1-α-hydroxylase enzyme plays an important role.

Vitamin D is fat-soluble vitamin, and is increasingly recognized to play a role in lipid metabolism: vitamin D deficiency is known to be highly-prevalent in obese people [32–37]and body fat content is negatively correlated with serum 25(OH)D levels [38–41]. We previously reported that vitamin D deficiency was associated with an increase in fatty infiltration of muscle in healthy young women [42] and showed that adequate dietary vitamin D prevents lipid accumulation in murine muscle [43]. At the molecular level, *in vitro* administration of 1,25(OH)_2_D to 3T3-L1 adipocytes caused significant inhibition of lipid accumulation and adipogenesis, with decreased lipid mobilization and availability [35]. To investigate the potential link between vitamin D and lipid metabolism, this study explores the role of tumoral CYP27B1 and local vitamin D synthesis in lipid changes detected in hyperplasia, early and late mammary carcinoma tissues and in the surrounding mesenchymal stroma.

Matrix-Assisted Laser Desorption Ionization Mass Spectrometry Imaging (MALDI MSI) is a powerful technique for the *in situ* analysis and imaging of the molecular composition of a broad spectrum of analytes such as proteins, peptides, drugs and lipids on tissue samples. This method is non-destructive and label-free, and allows multiplex analysis of hundreds to thousands of molecules simultaneously in the same tissue section [44,45]. The great advantage of MALDI MSI is the correlation of molecular information with traditional histology by keeping the spatial localization information of the analytes after mass spectrometric measurement. Frozen tissue sections are mounted on a metal plate coated with an ultra-violet (UV) absorbing matrix and placed in a mass spectrometer. A pulsed UV laser desorbs and analyzes analytes from the tissue, and their mass to charge (*m/z*) values are determined by a mass analyzer. Because the histological sample is divided into a raster of microsections covering the whole surface, molecular images are created of mass spectra from molecules present within each segment of the irradiated area [46]. The undamaged tissue sample can subsequently be stained with conventional methods such as hematoxylin and eosin (H&E) or immunohistochemistry, and co-registered to the mass spectrometric analysis. The mass signals (*m/z*) detected are then visualized as color intensity maps assigning molecular patterns to cell types. These color signals allow the detection of patterns, which represent the distribution in tissue of the molecules of interest [45]. This provides information on the identity and abundance of target molecules as well as their distribution within the tissue. Because analyte signals can be correlated with underlying tissue architecture without any geometrical distortion, MALDI MSI allows “molecular histology”. Consequently, pathologists are now able to correlate the distribution of specific compounds with pathologically interesting features [45,47–49].

As previously demonstrated, breast cancer is associated with alterations in breast lipid composition [50–55]. As summarized by Kawashima’s group, MALDI MSI has been widely used in cancer research for several reasons [56]. First, it is a powerful and versatile technique to directly visualize lipid distributions and relative abundance within tissue sections. Second, it allows researchers to detect molecules that despite being in very low abundance may be biologically important. Lipid profiles change in malignant epithelial regions allowing for distinguishment from surrounding stroma. As such, phospholipid signals could potentially serve both as biomarkers for classifying different subtypes of breast cancers as well as a biomarker of tumor progression. Other studies have corroborated the diagnostic ability of MALDI MSI on breast cancer [57] and its potential to identify subtypes of breast cancer with high efficiency and accuracy [58].

In the present study, using the innovative MALDI MSI technology combined with the MMTV-PyMT mouse mammary tumor model with and without *Cyp27b1* ablation, we demonstrate that mammary tumor-produced 1,25(OH)_2_D is mechanistically linked to lipid metabolism changes in hyperplasia and early & late carcinoma tissues. The scheme of the study design is presented in Suppl. Figure 1.

## Materials and Methods

### Conditional ablation of *Cyp27b1* in mouse

The MMTV-PyMT model of breast cancer displays highly aggressive spontaneous mammary tumors closely mimicking the human disease in its progression from premalignant to malignant stage followed by high metastatic frequency, and in its expression of poor outcome biomarkers.

*Cyp27b1^flox/flox^*mice were obtained with the gene *Cyp27b1* gene exon 8 floxed [59], and were further backcrossed with FVB mice (Friend leukemia Virus B-type,Charles River, Quebec) for a 99% FVB/NJ background. The mice colony was further bred with MMTV-PyMT transgenic mice (strain 634) and with MMTV-*Cre* mice on a FVB background, to produce several genotypes) including MMTV-PyMT; *Cyp27b1^flox/flox^*; *Cre*^+^ (homozygous), MMTV-PyMT ; *Cyp27b1^flox/flox^*; *Cre*^-^(control), and MMTV-PyMT ; *Cyp27b1^flox/+^*; *Cre*^+^ mice (heterozygous).

### Mammary tumor tissue extraction at specific development time points

A pattern of breast tumor development in the mammary glands was observed in our MMTV-PyMT animal model, in which the breast tumors initially appeared in left thoracic mammary glands then left abdominal mammary glands, then right thoracic mammary glands and finally right abdominal mammary glands. In this study, left thoracic mammary glands (primary tumors) and right abdominal mammary glands were collected from non-ablated and ablated mice at three developmental time points: hyperplasia (HP, week 5), early adenoma (MI, week 7), and carcinoma (LA, week 11) [60], embedded in optimal cutting temperature compound (ThermoFisher Scientific, USA), flash frozen and stored at -80°C until analyzed. Table 1 summarizes all the breast tumor samples collected and analyzed.

**Table 1.**
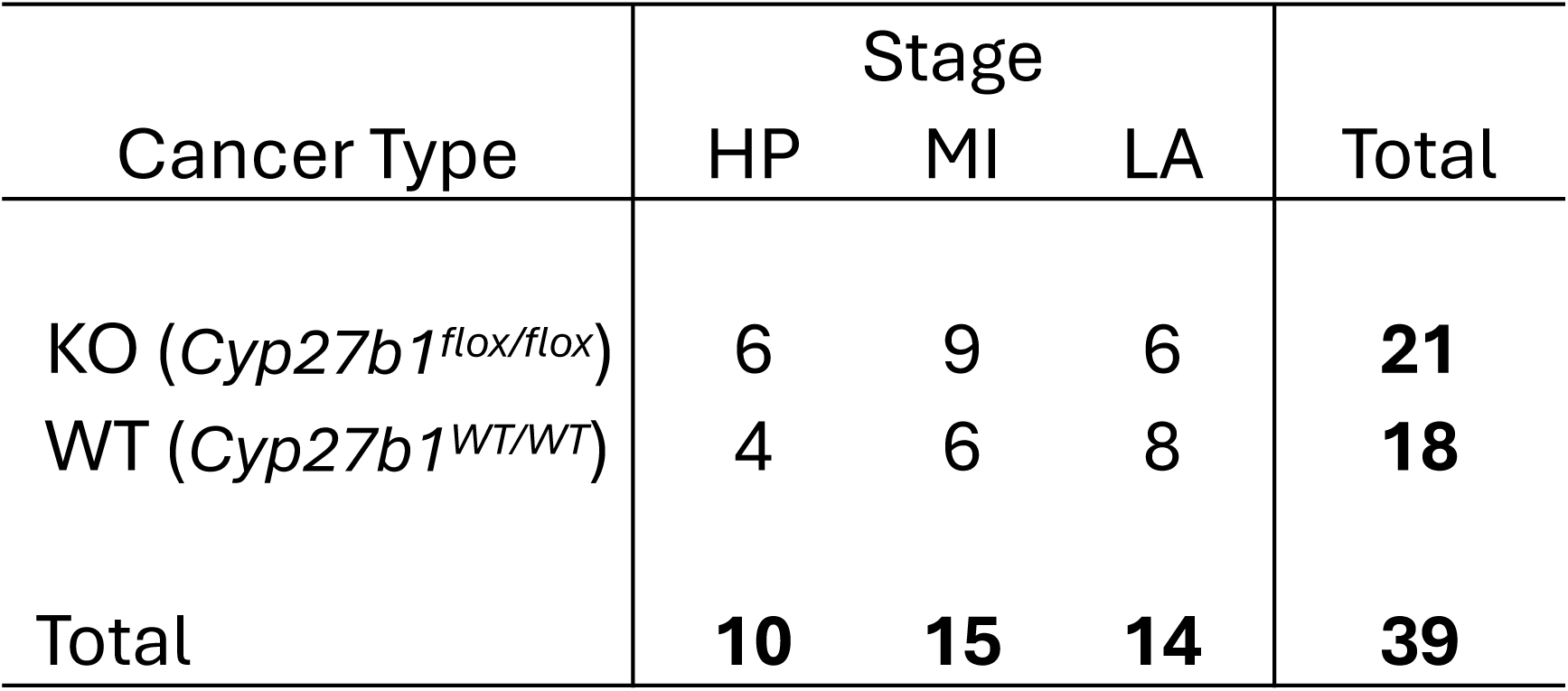
HP, hyperplasia (week 5) ; MS, middle stage (week 7) and LS, late stage (week 11) mammary tumors from *Cyp27b1^flox/flox^* (KO) and *Cyp27b1^WT/WT^* (WT) mice analyzed by MALDI MSI.

### Tissue sectioning and matrix sublimation

Flash frozen mammary tumors embedded in optimal cutting temperature compound were sectioned (16 μm thickness) at -25 °C with a ThermoScientific HM 560 cryostat. Sample sections were immediately thaw-mounted on indium-tin oxide-coated (ITO) glass slides (Delta Technologies ld., Loveland, CO, USA). Serial sections were obtained and stained with Haemotoxylin and Eosin (H&E) according to standard protocols. Liver homogenate sections were also deposited on ITO slides for systematic data normalization across slides and experiments. Sections were coated with the MALDI matrix 1,5-diaminoapthalene (DAN) in a home-built sublimation apparatus connected to a vacuum pump [61]. The optimal matrix thickness (corresponding to approximately 300 μg/cm^2^ of matrix) was obtained after 12 minutes of sublimation at 165°C and a vacuum pressure of 0.2 mbar.

### MALDI mass spectrometry

Dual polarity MALDI MSI data were acquired using a MALDI TOF/TOF (time of flight) ultrafleXtreme mass spectrometer (Bruker Daltonics, Billerica MA) in reflection geometry at an acceleration voltage of + or -20 kV, and laser repetition rate of 2 kHz (SmartBeam II Nd:Yag/355 nm laser with a focus of 15 μm in diameter for the minimum focus setting) on the entire section surface at a lateral spatial resolution of 100 μm. Positive and negative ionization mode data were acquired serially on the same tissue section as per protocol outlined in [61], with the positive grid array offset by 50 μm in both the x and y dimensions with respect to the negative data grid array. MSI data was acquired in the *m/z* 500-1000 range. In positive ionization mode, 150 laser shots were summed at each raster position, while 200 laser shots per array position were acquired in negative ionization mode. All other parameters, including laser fluency, delayed extraction and, detector gain were adjusted for the highest signal to noise ratio (S/N) and mass resolution in the *m/z* range under investigation. FlexImaging v4.1 was used for acquiring the MSI data (Bruker Daltonics, Billerica MA). Internal calibration was performed using known lipid peaks from the tissue providing a ∼10 ppm mass accuracy on lipid signals.

MALDI MS/MS data was acquired in LIFT mode using the ultrafleXtreme mass spectrometer. The parent ion mass selection window was set at +/-1 Da. Lipid identification was based on comparisons made with corresponding molecular weight entries in Lipidmaps (https://lipidmaps.org/), or with the existing lipid MALDI MSI literature.

### Data analysis

MSI data processing and analysis is schematized in Suppl. Figure 2. Raw data acquired from MALDI MSI was converted into the vendor-neutral imzML format after loading the data in FlexImaging v4.1 (Bruker Daltonik GmbH, Germany). Before conversion, the data was internally calibrated by flexAnalysis 3.4 Batch Process (Bruker Daltonik GmbH, Germany). The Cardinal package (v 1.4) in the R environment (v 3.4.3) was used for data pre-processing including baseline subtraction, peak picking, peak alignment and data reduction. After pre-processing, the reduced data from the positive and negative polarity of the same tissue sample were recombined into one imzML data file with an artificial *m/z* offset of 2000 for the negative polarity [62]. Unsupervised analysis of the MSI data was then performed using the spatially aware k-means segmentation in the Cardinal package to group pixels of similar characteristics with parameter setup of r=1, s=1, k=3. The number of segments was determined based on the tissue histology.

### Principal Component Analysis (PCA)

Principal component analysis (PCA) was performed on the lipid signals that highly correlated to three main clusters: hyperplasia, tumors, and mesenchymal stromal regions in the mammary tissues. Due to the high complexity of the reduced dataset, we chose to use PCA as the statistical method for data mining and visualization of those biological variants. In PCA, score images were created to represent the entire dataset and show prominent patterns with high intensity correlating to first principal components [63], and *m/z* values with highest loadings were also selected (analysis of variance (ANOVA) and Turkey test) to identify discriminatory lipid signals.

### Study approval

This animal study was approved by the McGill University Animal Compliance office. All procedures have followed an animal use protocol approved by Animal Care and Use Committee of McGill University (Protocol number 3520). This protocol has also followed ethical guidelines of the Canadian Council on Animal Care.

## Results

### MALDI MSI data clustering accurately identifies and differentiates tumor, mesenchymal stromal and adjacent muscle tissues in mammary glands of MMTV-PyMT mice

We previously generated a MMTV-PyMT mouse model using a Cre-loxP-mediated system [5,59] to obtain a homozygous ablation of gene *Cyp27b1* specific to the mammary epithelium (ME). The *Cyp27b1^flox/flox^*; *Cre*^+^ mice were ablated of vitamin D 1-α-hydroxylase in ME. In contrast, the *Cyp27b1^WT/WT^*; *Cre*^+^ mice expressed vitamin D 1-α-hydroxylase in ME and were considered as control animals (Cre^+^ WT). Lin *et al*. described malignancy progression in the mouse model as hyperplasia at 6 weeks, adenoma/mammary intraepithelial neoplasia (MIN) at 8 weeks, early carcinoma at 10 weeks and late carcinoma at 12 weeks [60]. Initial proof of concept measurements was performed on samples derived from these mouse models (i.e., control: *Cyp27b1^WT/WT^*; *Cre*^+^ & experimental: *Cyp27b1^flox/flox^*; *Cre*^+^) MALDI MSI analyses were performed on *Cyp27b1*-ablated and *Cre*^+^ WT tumor specimens for each of these developmental time points (Figure 1). Histological evaluation demonstrated that for both genotypes, a rapid progression in tumor development was observed with time (Figure 1A). However, at 6 weeks, tumor cell differentiation appeared to be more advanced in *Cyp27b1*-ablated breast tissue than in wild type mice. This finding agrees with our previous observation in which hyperplasia displays earlier appearance in MMTV-PyMT mice following the ablation of the *Cyp27b1* gene in mammary epithelium [5]. More areas of solid tumors were observed at weeks 8-10 in *Cyp27b1*-ablated mice compared to wild type mice. Increased enlargement in tumoral area and reduction in stromal area are displayed at 12 weeks in both *Cyp27b1*-ablated tumors and wild type tumors. MSI results confirmed the more rapid tumor progression in *Cyp27b1*-ablated mice compared to wild type mice. This is exemplified when monitoring the spatial distribution (Figure 1B) and relative abundance (Figure 1C) or selected lipid signals at *m/z* 722.54 and *m/z* 734.60 clearly associated with tumor histology. Indeed, the intensity of both signals strongly increases in tumor areas as a function of time. We however observed that the relative abundance of these signals was always associated with a lower yield in the *Cre*^+^ WT versus the *Cyp27b1*-ablated tissue samples, again confirming the faster development of cancer in the *Cyp27b1*-ablated versus the *Cre*^+^ WT mice.

**Figure 1.**
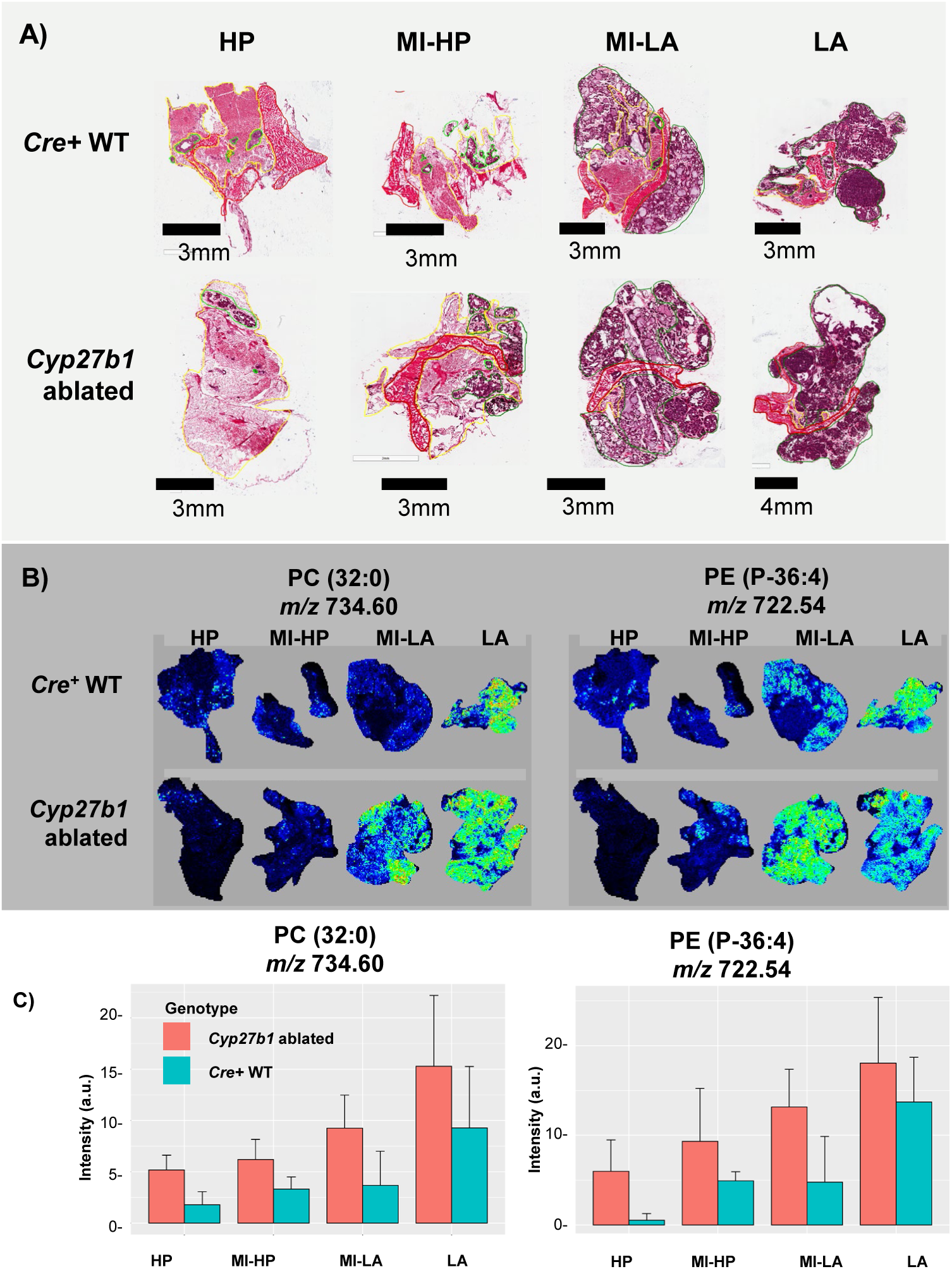
Histology and MALDI MSI of hyperplastic and tumoral cancer lesions from breast tumor tissue of Cre^+^ WT and *Cyp27b1-*ablated MMTV-PyMT mice at 6 weeks (hyperplasia, HP), at 8 weeks (adenoma/mammary intraepithelial neoplasia, MI-HP), 10 weeks (early carcinoma, MI-LA) and 12 weeks (late carcinoma, LA). A) H&E staining demonstrating a cancer progression from hyperplasia to carcinoma with time. B) MALDI MSI density maps for two selected lipid ion signals (*m/z* 734.60 and 722.54) strongly associated with carcinoma histology. C) Lipid signal ion intensity as a function of time and sample type demonstrating a faster cancer progression in the *Cyp27b1-*ablated mice.

For all further histology and MALDI MSI measurements, we re-defined malignancy stages as hyperplasia (HP) at 5 weeks, middle stage (MI) at 7 weeks and late stage (LA) at 11 weeks (Table 1). A cohort consisting of 10 HP (week 5), 15 MI (week 7) and 14 LA (week 11) breast tumor samples were analyzed. Clustering is a powerful tool to group pixels from MALDI MSI with similar mass spectra. Here, structural lipid clusters were generated in a completely unsupervised manner without previous knowledge of anatomical structures of the tissues. Crucially, clustering of MALDI MSI images displayed a high correspondence in lipid profiles to hematoxylin/eosin areas for hyperplasia/tumor, mesenchymal stroma and adjacent muscle tissue in both *Cyp27b1*-ablated and control mice at all time points. Based on MALDI MSI results (Figure 2), accurate overlays between histology and the data segments were observed. Clustering of MALDI MSI can therefore allow to pinpoint lipid expressions in particular anatomical structures or histologies. Further, a clear progression of cancer was observed with a progressive appearance of a data segment associated with hyperplasia/tumor confirming our initial measurements. A more detailed view of the hyperplasia/tumor, mesenchymal stroma segmentation can also be found in Suppl. Figure 1.

**Figure 2.**
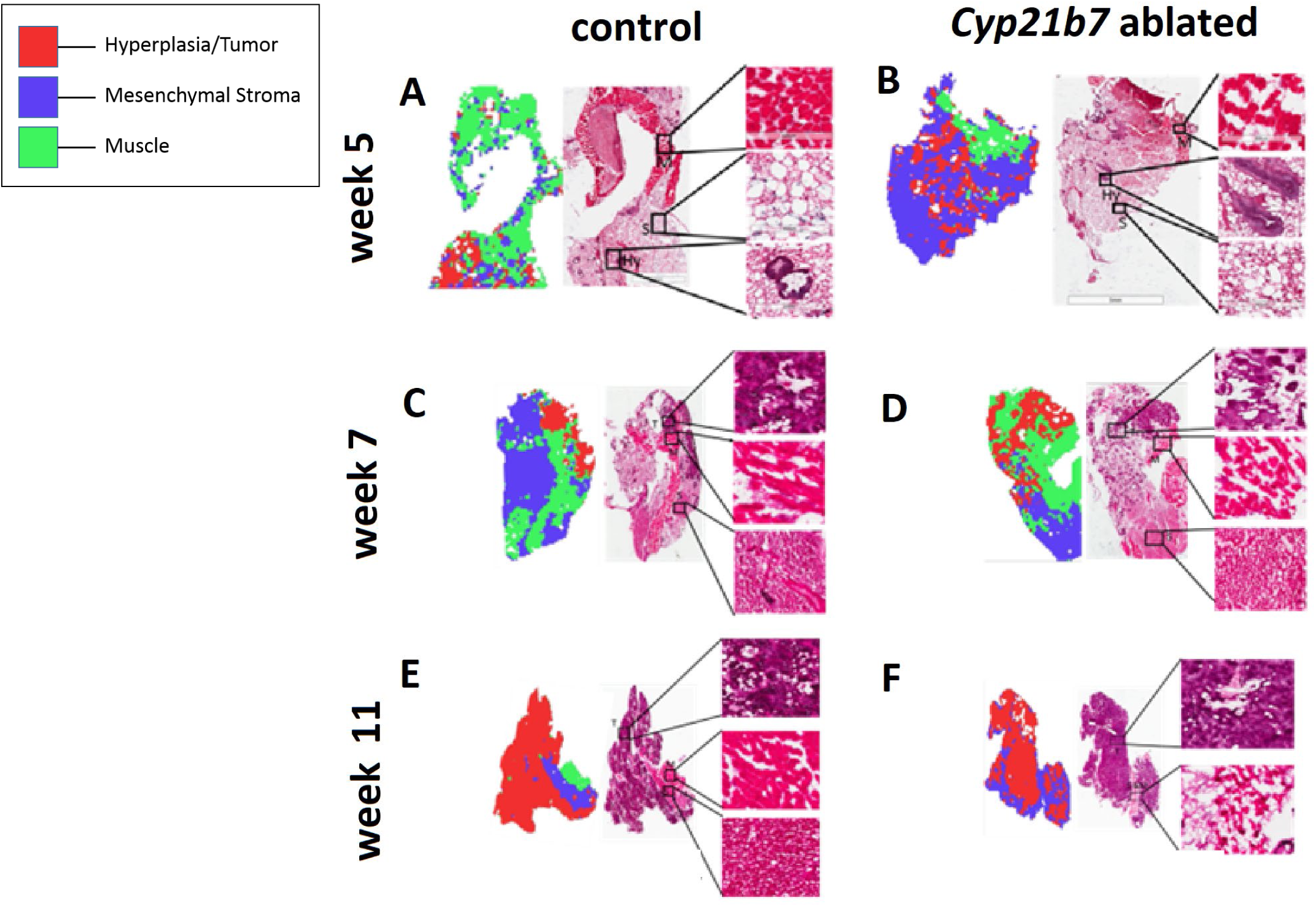
H&E staining (right) and MALDI MSI data segmentation (left) of WT (A,C,E) and *Cyp27b1-*ablated MMTV-PyMT mice (B,D,F) at 5, 7 and 11 weeks of mammary gland tissue development. The segments allow to differentiate hyperplasia/tumor, mesenchymal stroma and muscle histologies. Throughout the 11-week period, hyperplasia/tumor areas were observed to increase in surface while the mesenchymal stroma areas were reduced.

### Co-existence of hyperplastic and tumoral lesion is observed in mammary glands of MMTV-PyMT mice at 7 weeks with and without *Cyp27b1* ablation

Tumor heterogeneity causes morphological, genetic and behavioral variability within a single breast tumor [64]. When tumoral MALDI MSI regions results were further analysed, two distinct segments corresponding to hyperplastic and tumoral cancer lesions could be identified. Again, lipid profiles indicated a time-wise expression progression from week 5 (all hyperplasia) through 7 (mix) to 11 (all tumor) in both the non-ablated and *Cyp27b1*-ablated tumors (Figure 3 A,B,C and D,E,F respectively). The two distinct lipid profile sub-segments present at week 7 show accurate correspondence to histological structures of hyperplastic and tumoral lesions, indicating differential lipid expressions for hyperplasia and tumor within the same mammary tissue in both control (Figure 3B) and *Cyp27b1*-ablated (Figure 3E) animals. These results suggest that in addition to progression in cell morphology, mammary tumors display heterogeneity in lipid expression patterns in tumor cells.

**Figure 3.**
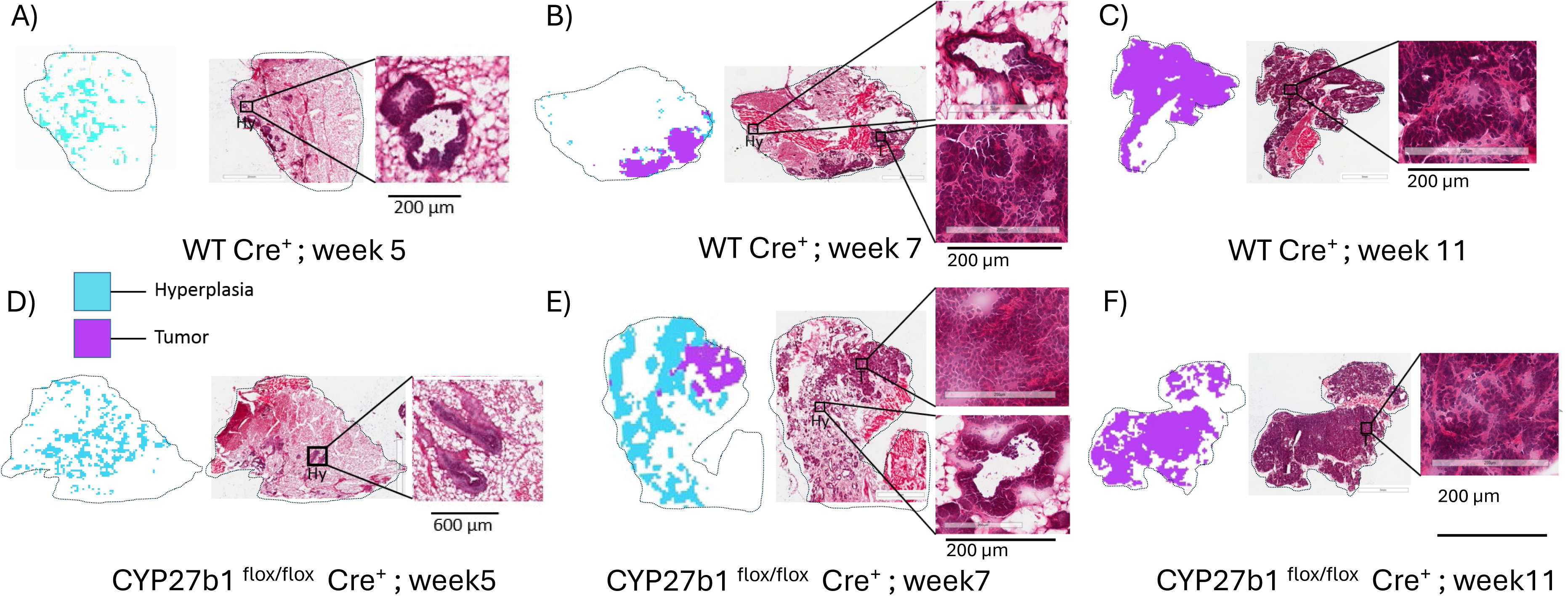
H&E staining (right) and MALDI MSI data segmentation (left) differentiating hyperplastic and tumoral cancer lesions from breast tumor tissue of WT (A, B, C) and *Cyp27b1-*ablated MMTV-PyMT (D, E, F) mice at 5, 7 and 11 weeks of mammary gland tissue development. At week 5, only hyperplasia was observed in both Cre+ WT tissue and Cyp27b1-ablated breasts tissue (A, D). At 7 weeks, the co-existence of hyperplastic and tumoral lesions is observed. Segmentation of MALDI MSI showed differential lipid expression in hyperplasia and tumor regions within the same mammary tissue in both WT and ablated tissue (B, E). At 11 weeks, segmentation exclusively showed tumoral lipid profiles in tumor regions corresponding to the fully differentiated mammary tumor cells as observed by H&E staining (C, F).

### Lipid composition changes during mammary tumor development

To determine main features that differentiated the histological regions and stages, we ran a global PCA analysis on all samples to determine the closest correlation with defined independent variables such as development stage, genotype or histology (Figure 4). PCA analyses revealed that the strongest percentage of explained variation was found across the different cancer development stages from hyperplasia to carcinoma. It also allowed to remove undesired features to identify the top lipid markers by ANOVA.

**Figure 4.**
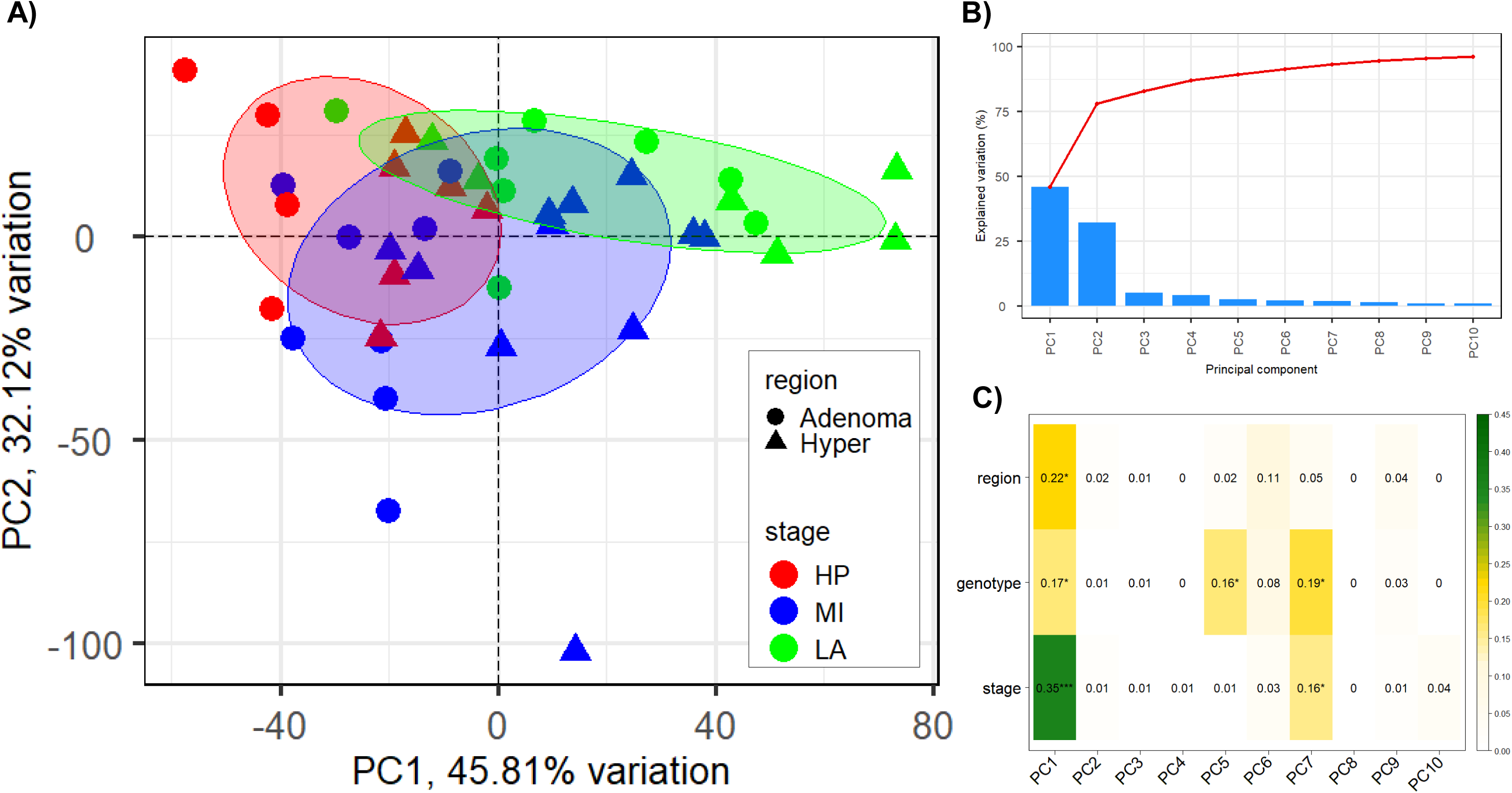
A) PCA biplot analysis of the hyperplastic and tumoral cancer lesions from breast tumor tissue of WT and *Cyp27b1-*ablated MMTV-PyMT mice at 5 (HP), 7 (MI) and 11 (LA) weeks of mammary gland tissue development. B) Scree plot of the top ten components. C) Pearson correlation of top 10 components. The top discriminant features are stage, region and genotype.

Top *m/z* features obtained from PCA analysis (Tukey test after ANOVA, P values < 0.05) showed significant progressive alterations throughout mammary tumor progression (Suppl. Figure 4). MALDI MS/MS tandem mass spectrometry performed directly on the tissue sections was conducted to identify the subtypes of these molecular signatures based on their corresponding *m/z* ratio in control animals and *Cyp27b1*-ablated animals (Suppl. Table 1). Interestingly, most of the top 16 discriminating lipid ion signals were observed in the negative ion mode. All of these lipid signals were up-regulated throughout mammary tumor progression. For the control and *Cyp27b1*-ablated animals, more detailed independent analyses were performed to identify the top lipid signals associated with cancer progression (Figure 5). In this case, the corresponding MALDI MSI datasets were used towards PCA analyses. Out of the 9 and 11 phospholipid signals associated with cancer progression in the control and *Cyp27b1*-ablated animals respectively, 8 are held in common. These belong to several phospholipid families including phosphocholines, phosphoethanolamines, and phosphatidic acids. Other comparisons were performed across the different cancer progression stages for the control and *Cyp27b1*-ablated samples. We specifically compared: HP week 5 to HP week 7, HP week 7 to Tumor week 7, and Tumor week 7 to Tumor week 11. Surprisingly, even if histology showed significant differences, no consistent lipid expression differences were found when comparing hyperplasia and tumor at week 7. Only when comparing tumors between weeks 7 and 11 could lipid signals be associated with cancer progression (Suppl. Figure 5). For each sample class, 6 lipid markers were found differentially expressed, only 3 of which were held in common. These results show that consistent differences in lipid abundance can be observed between the control and the *Cyp27b1*-ablated tumors. Because cancer progression was observed at a faster rate for the *Cyp27b1*-ablated animals, we also compared hyperplasia in samples from the *Cyp27b1*-ablated at week 5 to the control at week 7. However, no significant differences were observed. Finally, we compared control and *Cyp27b1*-ablated tumor areas at week 11 and found 2 differentially expressed lipid signals further emphasising that consistent molecular differences exist between the control and *Cyp27b1*-ablated tumors (Suppl. Figure 6).

**Figure 5.**
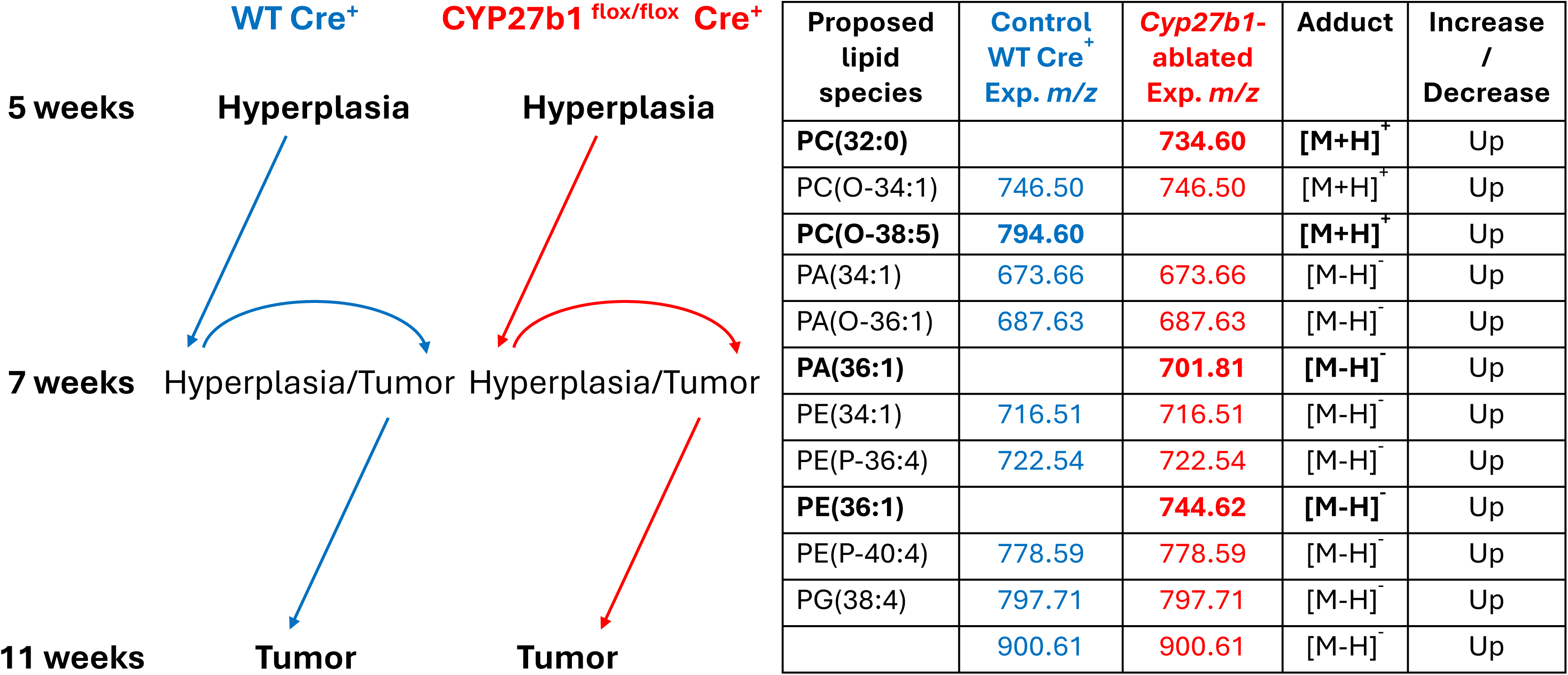
Top *m/z* features obtained from PCA analysis (Tukey test after ANOVA, P values < 0.05) showing significant progressive alterations throughout mammary tumor progression for the WT Cre^+^ and the CYP27b1 ^flox/flox^ Cre^+^ mice.

## Discussion

Vitamin D has been investigated for its protective effect against breast cancer in numerous studies, but evidence including meta-analysis regarding circulating and supplemental vitamin D on breast cancer risk remains inconclusive [29,65,66]. Nonetheless, low circulating 25(OH)D concentration in breast cancer patients correlates with increased overall mortality and distant recurrence [30,67].

Evidence for lipid alteration leading to breast cancer progression is suggested by compositional change in lipid droplets observed in MCF7 and MDA-MB-231 malignant breast cancer cells [68]. Furthermore, 1,25(OH)_2_D exhibits regulatory effect on adipogenesis through modifying transcriptional factors C/EBP and PPAR-γ in 3T3-L1 preadipocytes [69]. Thus, locally produced 1-α-hydroxylase (encoded by the *Cyp27b1*gene) which mediates conversion of the prohormone 25(OH)D to its bioactive form 1,25(OH)_2_D [70,71] may play a crucial role on lipid metabolism-linked breast tumor progression.

To examine the overall lipid profiles in primary breast tumors in MMTV-PyMT mice, data segmentation was performed on the information acquired by MALDI MSI in an unsupervised manner. We show that individual segments accurately reflect the anatomical structures of breast tumor tissue (Figure 2), suggesting differential lipid compositions in tumors, mesenchymal stroma and adjacent muscle tissues. Additionally, adjacent hyperplastic and tumoral regions from primary breast tumors of MMTV-PyMT mice displayed differential lipid expressions at 7 weeks (Figure 3B and E), which highlights that breast tumor exhibits intratumoral heterogeneity from various subpopulations of tumor cells [72]. It is also remarkable that alterations in lipid composition are seen in normal-appearing tissues without visible histopathological alterations adjacent to hyperplastic lesions in both Cre^+^ WT (Figure 3A) and *Cyp27b1*-ablated breast tissues (Figure 3D).This crucial observation reflects the high sensitivity of lipid profiling in breast tumor progression prior to histopathological changes, and this finding agrees with previous observations of the strong association between tumor grade and tissue phospholipid profiles in human breast cancer biopsies [73].

A potential link between the abundance of PC (phosphatidylcholine), PE (phosphatidylethanolamine) and PA (phosphatidic acids) in tumor and breast cancer malignancy during late carcinoma stage (Figure 5) was also observed. To be more specific, out of 16 up-regulated lipid biomarkers throughout breast tumor neoplasia, the majority (12/16) consists of 4 PC (3 of which are alkyl ether substituted), 6 PE and 2 PA (Supplemental Figure 4). These findings are consistent with the study from Eiriksson et al., (2020) who found elevated levels of PCs and PEs in some breast cancer cell lines [74]. In a cell line representing HER2-overexpressing tumor subtype, an elevated expression of PC and PE containing short-chained (≤ C-16) saturated or monounsaturated fatty acids were observed. Further, increased abundance of PC ≥ C-40 was found in cell lines of triple negative BC subtype [74]. Hosokawa et al., (2017) also found that PC(31:1) was associated with a higher recurrence of breast cancer after the analysis of human tumors by MALDI MSI [75]. Finally, Tan et al. (2020) identified PAs in patient’s serum as potential markers of breast cancer progression [76].

Our previous study showed an acceleration in hyperplasia appeared in the mammary glands with ablation of tumor-synthesized 1,25(OH)_2_D in MMTV-PyMT mouse model of breast cancer [77]. To investigate the direct effect of 1,25(OH)_2_D on lipid compositions in breast tumors, it is important to compare primary breast tumors at the same tumoral progression stage. According to PCA analysis, *CYP27b1* ablation causes differential lipid expression in hyperplastic areas (Figure 4), suggesting a modifying effect of locally produced 1,25(OH)_2_D on lipid metabolism during the hyperplasia stage of breast tumor development. This finding is also supported by the significant difference in abundance of top lipids markers in these hyperplastic regions, including lipid *m/z* 794.60 which was exclusively expressed in control mice (Figure 5).

Two potential mechanisms can be proposed based on our observations. First, tumor-produced 1,25(OH)_2_D may directly affect lipid expression in tumor cells which then regulates breast tumor progression (Figure 6A, pathway 1). Thus, the lack of tumoral 1,25(OH)_2_D production may lead to phospholipid accumulation in breast tumor cells, accelerating tumor progression (Figure 6B, pathway 1). Alternatively, the accelerated development of hyperplasia observed in *Cyp27b1*-ablated tumors [5] may be responsible for the lipid compositional changes within tumor cells (Figure 6A, pathway 2). Supporting the hypothesised mechanism for a direct effect of tumor produced 1,25(OH)_2_D on lipid expression is the observation that changes in lipid composition are already apparent in histologically normal mammary tissue, before any evidence of hyperplasia. Our previous observations of 1,25(OH)_2_D’s dowregulation of PPAR-γ (peroxisome proliferator-activated receptor gamma), a transcriptional factor required for adipogenesis [78] in bone marow of SAM-P/6 mice [79] suggests a possible association of 1,25(OH)_2_D and lipid metabolism by modification of lipid-related gene expression in breast tumor cells (Figure 6A, pathway 3). Thus, we suspect that ablation of tumor produced 1,25(OH)_2_D causes altered expression of the genes involved in lipid metabolism, affecting breast tumor cells (Figure 6B, pathway 3). Because VDR is expessed in both tumor cells and normal adjacent stromal cells in human breast cancer tissue [80] it is tempting to speculate that lipid composition in the mesenchymal stroma may be regulated by tumor-released 1,25(OH)_2_D acting through VDR expressed on adjacent stromal cells (Figure 6A, pathway 4).Further studies are needed to examine this potential mechanism.

**Figure 6.**
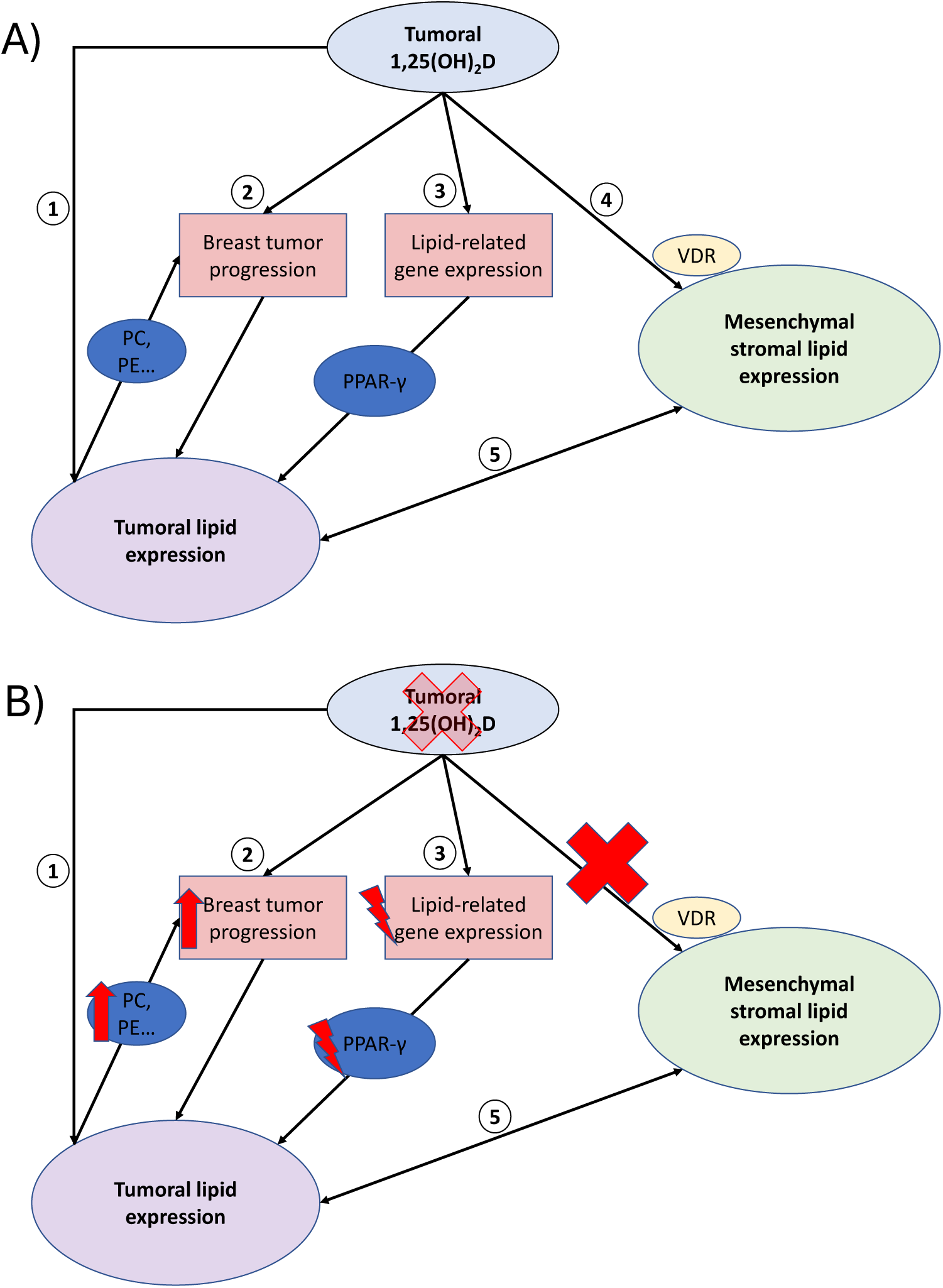
Potential mechanisms of association among tumoral 1,25(OH)_2_D, tumoral lipid expression and msenchymal stromal lipid expression in Cre^+^ WT tissues (A) and in Cyp27b1-ablated tissues (B). Pathway 1, tumoral produced 1,25(OH)_2_D alterates lipid expression in tumors, which results in breast tumor progression in Cre^+^ WT animals in A, CYP27B1 ablation leads to phospholipid accumulation such as phosphatidylcholine (PC), phosphatidylethanolamine (PE) and phosphatidic acids (PA) in tumor, resulting in accelerated breast tumor progression in (B); Pathway 2, tumoral lipid compositional change is a consequence of the regulation of tumoral 1,25(OH)_2_D on breast tumor progression (A), which ablation of CYP27b1 in mammary epithelium leads to an accelerated breast tumor progression and alterated tumoral lipid expression; Pathway 3, 1,25(OH)_2_D can regulate lipid-related gene expression such as in PPAR-γ to inhibit adipogenesis in breast tumors (A). Thus, the lack of tumoral produced 1,25(OH)_2_D may affect lipid-related gene expression (B); Pathway 4, tumoral produced 1,25(OH)_2_D regulates mesenchymal stromal lipid expression through vitamin D receptor (VDR) on stromal cells (A), which is inhibited under the condition of CYP27B1 ablation (B); Pathway 5, alteration in lipid profiles from mesenchymal stroma may be due to tumor-stromal interaction (A and B).

In conclusion, ablation of tumoral *CYP27b1* in a model of breast tumor progression reveals a novel role for 1,25(OH)_2_D as a lipid regulating hormone with inhibitory effcts on cancer progression. This supports previous studies that suggested an important association between vitamin D metabolism and breast tumor progression [5,6,81] Using our model of breast tumor progression, we show that lipid compositional change is an early event in breast cancer progression prior to any visible neoplastic change in breast tumors. Importantly, absence of tumoral CYP27B1 alters lipid compositions in hyperplastic lesions suggesting a mechanistic link between tumoral 1,25(OH)_2_D production, local lipid metabolism and breast tumor progression.

## Acknowledgements

The original *Cyp27b1 ^flox/flox^*mice were obtained from Dr René St-Arnaud(Genetics Unit, Shriners Hospital for Children, Montréal, QC, Canada, H3G 1A6).We thank Patrica Hales for excellent secretarial assistance.

## Notes

The authors have nothing to declare. This work was supported by grant number MOP 10839 from the Canadian Institutes of Health Research (CIHR) to RK. PC acknowledges funding from the Natural Sciences and Engineering Research Council of Canada (NSERC) for Discovery grant RGPIN/03125-2021. EY and RSP acknowledge funding from NSERC Postgraduate Scholarship-Doctoral program.

The authors declare no conflict of interest

### Competing Interest Statement

The authors have declared no competing interest.

