## supplemental figures for "Mass spectrometry imaging shows specific lipids signals linked to Cyp27b1 ablation in the MMTV PyMT breast cancer model"

| Cancer Type | Stage HP | Stage MI | Stage LA | Total |
|---|---|---|---|---|
| KO (*Cyp27b1*$^{flox/flox}$) | 6 | 9 | 6 | **21** |
| WT (*Cyp27b1*$^{WT/WT}$) | 4 | 6 | 8 | **18** |
| Total | **10** | **15** | **14** | **39** |

HP, Early Stage – Week 5 (Hyperplasia)

MI, Middle Stage – Week 7 (Mixed Cancer)

LA, Late Stage – Week 11 (Adenoma)

### Sample Preparation & Analysis

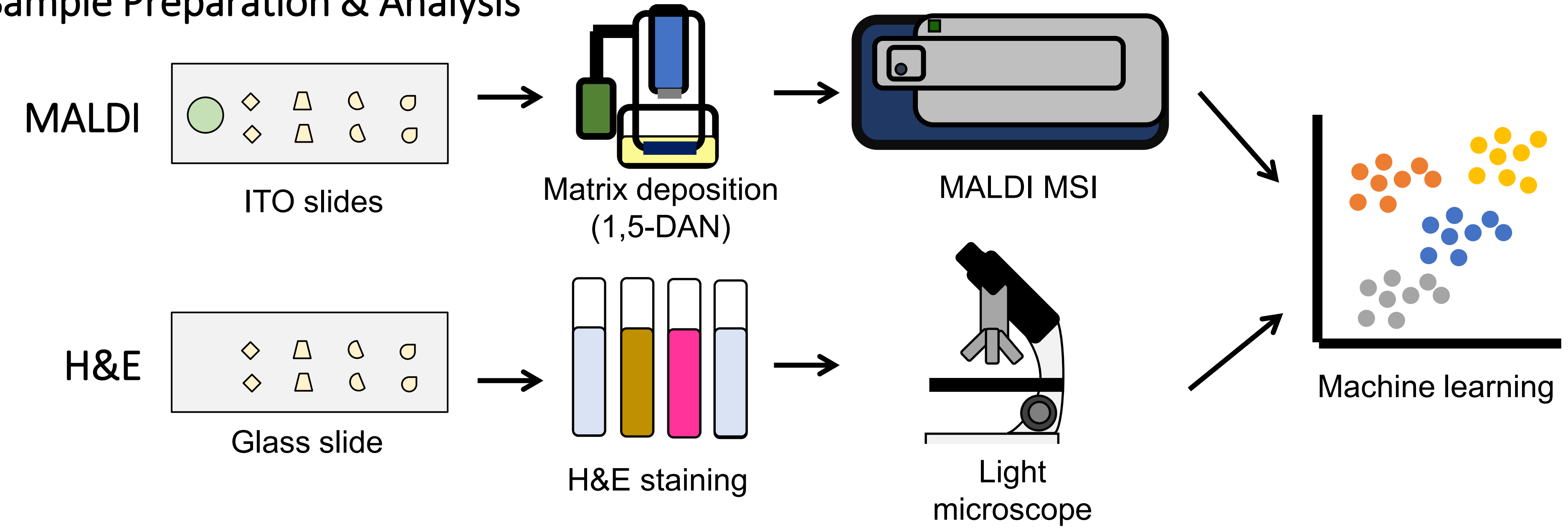


Suppl. Figure 1: Study design

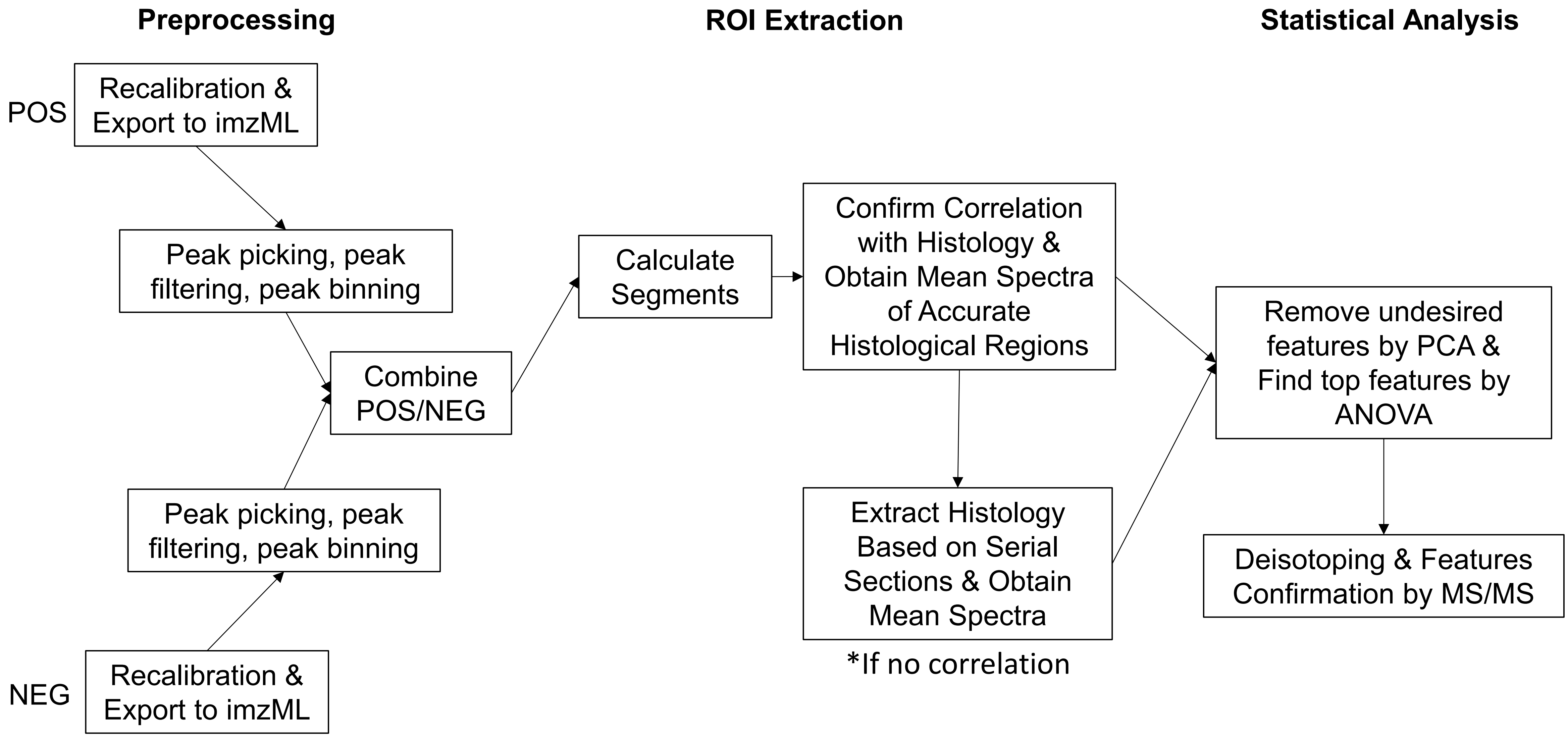


Suppl. Figure 2: MALDI MSI data processing pipeline.

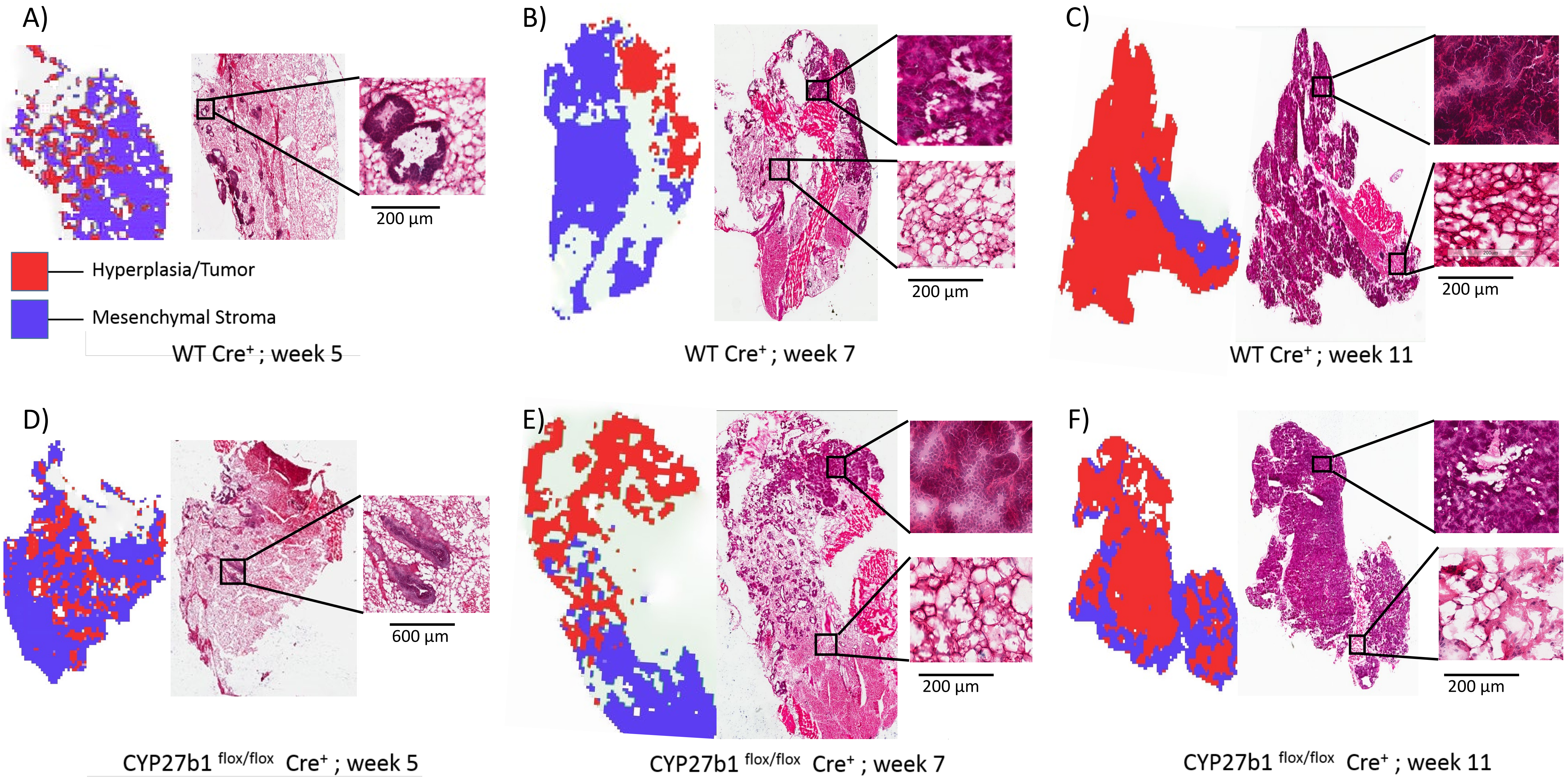


Suppl. Figure 3

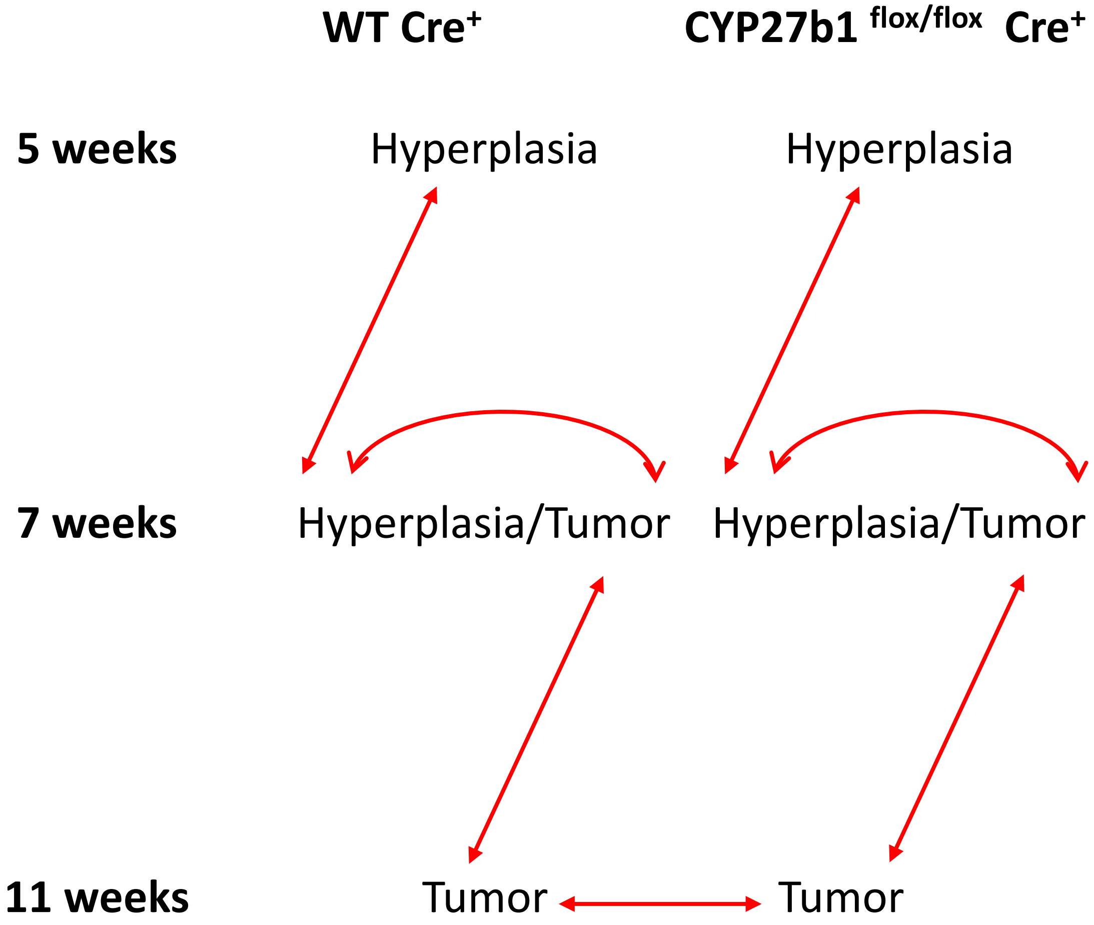


| Proposed Species | Experimental *m/z* | Adduct |
|---|---|---|
| PC(32:0) | 734.60 | $[M+H]^+$ |
| PC(O-34:1) | 746.50 | $[M+H]^+$ |
| PC(O-38:6) | 792.60 | $[M+H]^+$ |
| PC(O-38:5) | 794.60 | $[M+H]^+$ |
| | 642.67 | $[M-H]^-$ |
| PA(34:1) | 673.66 | $[M-H]^-$ |
| SM(d18:1/16:0) | 687.63 | $[M-CH_3]^-$ |
| PA(36:1) | 701.81 | $[M-H]^-$ |
| PE(34:1) | 716.51 | $[M-H]^-$ |
| PE(P-36:4) | 722.54 | $[M-H]^-$ |
| PE(136:1) | 744.62 | $[M-H]^-$ |
| PE(O-38:6) | 748.59 | $[M-H]^-$ |
| PE(P-38:4) | 750.58 | $[M-H]^-$ |
| PE(P-40:4) | 778.59 | $[M-H]^-$ |
| PG(38:4) | 797.71 | $[M-H]^-$ |
| | 900.61 | $[M-H]^-$ |

Suppl. Figure 4. Top *m/z* features obtained from PCA analysis (Tukey test after ANOVA, P values < 0.05) showing significant progressive alterations throughout mammary tumor progression for the WT Cre$^+$ and the CYP27b1 $^{flox/flox}$ Cre$^+$ mice, as well as between late-stage tumors.

| Experimental *m/z* | Theoretical *m/z* | Progression | Proposed Species | Adduct | Diagnostic Fragment Ions | Confirmation |
|---|---|---|---|---|---|---|
| 734.60 | 734.57 | Up | PC(16:0/16:0) | $[M+H]^+$ | (239), 184 | Berry et al. |
| 746.50 | 746.60 | Up | PC(O-16:0/18:1) | $[M+H]^+$ | (239), 184 | Berry et al. |
| 792.60 | 792.59 | Up | PC(O-38:6) | $[M+H]^+$ | 184 | Berry et al. |
| 794.60 | 794.61 | Up | PC(O-38:5) | $[M+H]^+$ | 184 | Berry et al. |
| 716.51 | 716.52 | Up | PE(16:0/18:1) | $[M-H]^-$ | 478, 460, 452, 434, 281, 255, 153, 140, 79 | Lipids Maps |
| 722.54 | 722.51 | Up | PE(P-16:0/20:4) | $[M-H]^-$ | 436, 303, 259,(239), (196), 153, 140, 97, 79 | Lipids Maps |
| 744.55 | 744.55 | Up | PE(18:0/18:1) | $[M-H]^-$ | 480, 462, 281, 153, 140, 97, 79 | Lipids Maps |
| 701.55 | 701.51 | | PA(18:0/18:1) | $[M-H]^-$ | 435, 417, 283, 153, 97, 79 | Lipids Maps |
| 687.57 | 687.53 | Up | SM(d18:1/16:0) | $[M-CH_3]^-$ | 281, 153, 79 | Lipids Maps |
| 673.66 | 673.48 | Up | PA(16:0/18:1) | $[M-H]^-$ | 435, 417, 409, 391, 281, 255, 153, 97, 79 | Lipids Maps |
| **642.52** | | Up | | $[M-H]^-$ | 79, 153, 255, 309, 392, 466, 486 | |
| **900.67** | | Up | | $[M-H]^-$ | 281, 153, 97, 79 | |
| 797.51 | 797.53 | Up | PG(18:0/20:4) | $[M-H]^-$ | 723, 437, 419, 303, 283, 153, 79 | Lipids Maps |
| 778.59 | 778.58 | Up | PE(P-20:0/20:4) | $[M-H]^-$ | 464, 446, 331, 153, 140, 97, 79 | Lipids Maps |
| 750.58 | 750.54 | Up | PE(P-18:0/20:4) | $[M-H]^-$ | 464, 446, 303, 259, 196, 153, 140, 97, 79 | Lipids Maps |
| 748.59 | 748.53 | Up | PE(O-16:0/22:6) | $[M-H]^-$ | 462, 444, 303, 259, 153, 140, 79 | Lipids Maps |

Suppl. Table 1. Identification of the 16 top discriminatory lipid signals showing significant progressive alterations throughout mammary tumor progression for the WT $Cre^+$ and the CYP27b1 $^{flox/flox}$ $Cre^+$ mice

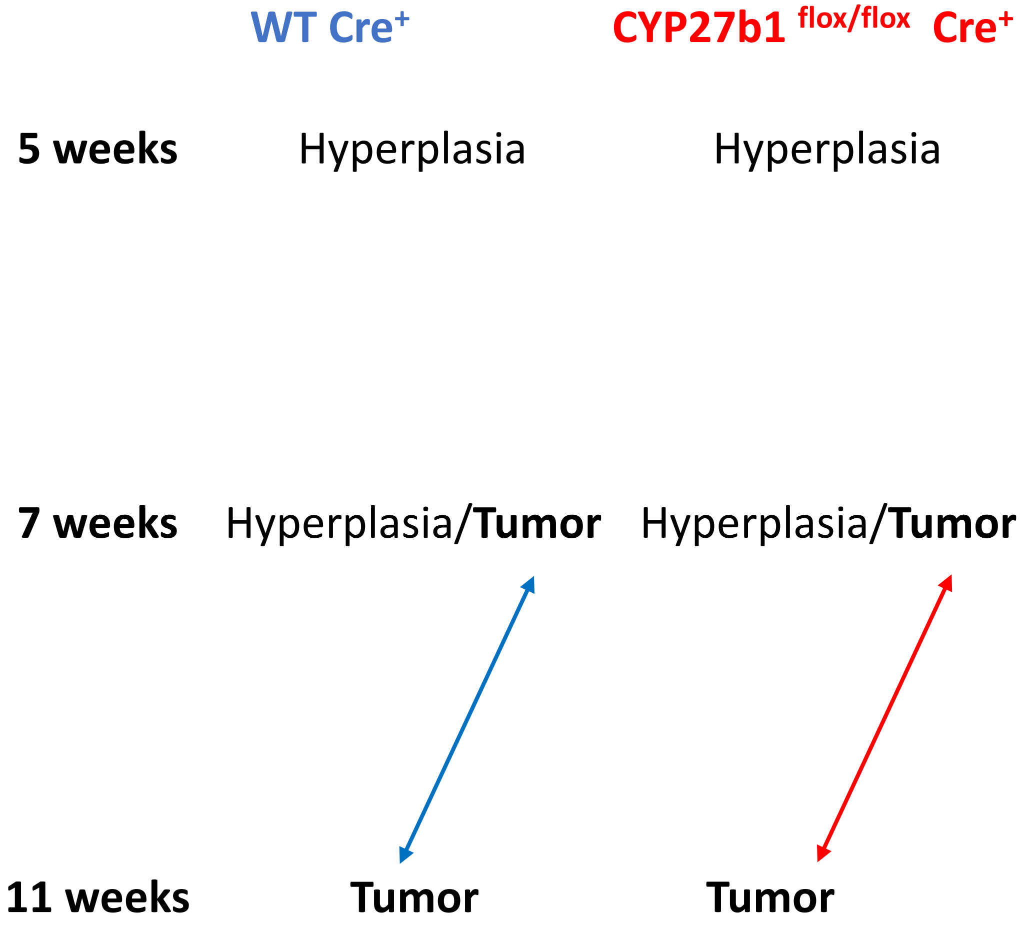


**WT Cre+**

| Proposed Species | Experimental *m/z* | Adduct |
|---|---|---|
| PC(O-34:1) | 746.50 | $[M+H]^+$ |
| PC(O-38:5) | 794.60 | $[M+H]^+$ |
| PA(16:0/18:1) | 673.66 | $[M-H]^-$ |
| **SM(d18:1/16:0)** | **687.63** | $[M-CH_3]^-$ |
| PE(P-20:0/20:4) | 778.59 | $[M-H]^-$ |
| **PG(18:0/20:4)** | **797.71** | $[M-H]^-$ |

**CYP27b1 flox/flox Cre+**

| Proposed Species | Experimental *m/z* | Adduct |
|---|---|---|
| PC(O-38:6) | 792.60 | $[M+H]^+$ |
| **SM(d18:1/16:0)** | **687.63** | $[M-CH_3]^-$ |
| PA(18:0/18:1) | 701.81 | $[M-H]^-$ |
| PE(16:0/18:1) | 716.51 | $[M-H]^-$ |
| **PG(18:0/20:4)** | **797.71** | $[M-H]^-$ |

Suppl. Figure 5. Top *m/z* features obtained from PCA analysis (Tukey test after ANOVA, P values < 0.05) showing significant progressive alterations for mammary tumor progression for the WT Cre+ and the CYP27b1 flox/flox Cre+ mice between week 7 and week 11.

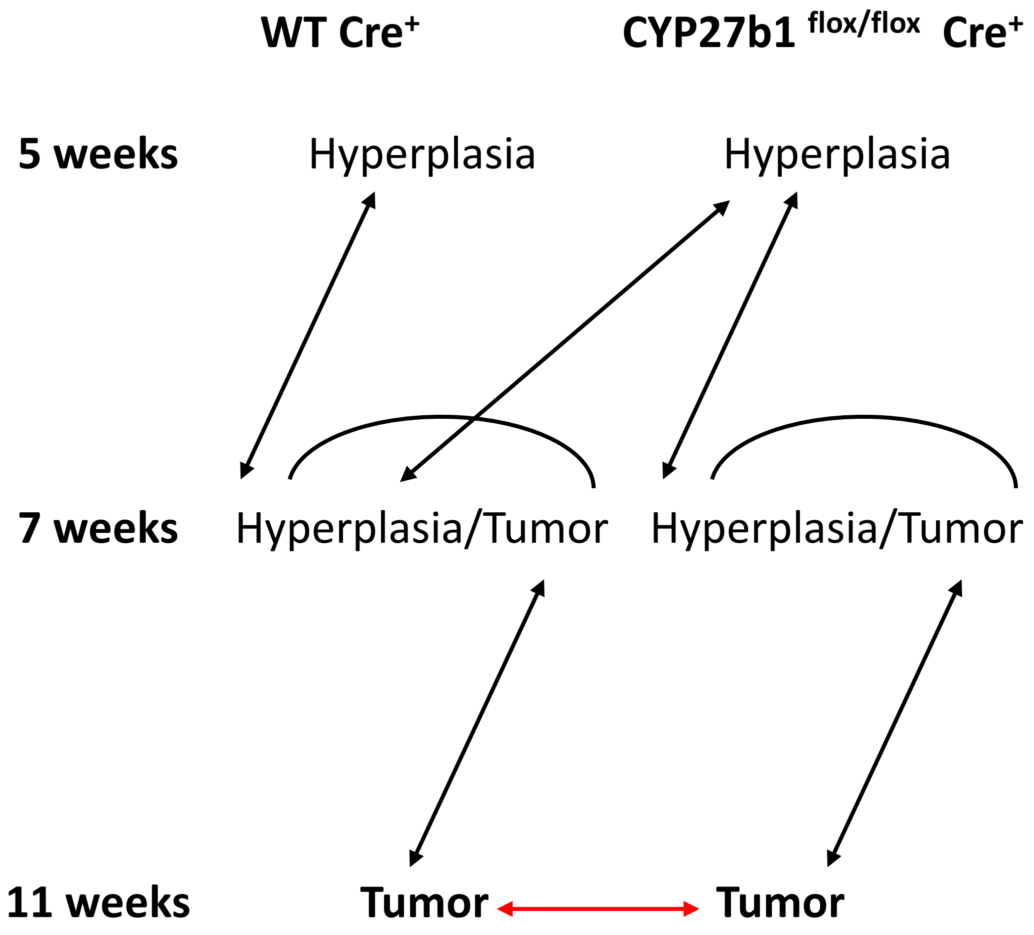


| Proposed Species | Experimental *m/z* | Adduct |
|---|---|---|
| SM(d18:1/16:0) | 687.63 | $[M\text{-}CH_3]^-$ |
| PA(18:0/18:1) | 701.81 | $[M\text{-}H]^-$ |

Suppl. Figure 6. Top *m/z* features obtained from PCA analysis (Tukey test after ANOVA, P values < 0.05) showing significant expression differences for week 11 mammary tumor for the WT Cre$^+$ and the CYP27b1 $^{flox/flox}$ Cre$^+$ mice.
